# Structural Dynamics of DNA Depending on Methylation Pattern

**DOI:** 10.1101/2020.06.29.178608

**Authors:** Takeru Kameda, Miho M. Suzuki, Akinori Awazu, Yuichi Togashi

**Affiliations:** Graduate School of Science, Hiroshima University, Higashi-Hiroshima, Hiroshima, Japan; RIKEN Center for Biosystems Dynamics Research (BDR), Wako, Saitama, Japan; Graduate School of Medicine, Nagoya University, Nagoya, Aichi, Japan; Graduate School of Integrated Sciences for Life, Hiroshima University, Higashi-Hiroshima, Hiroshima, Japan; RIKEN Center for Biosystems Dynamics Research (BDR), Higashi-Hiroshima, Hiroshima, Japan

## Abstract

DNA methylation is associated with a number of biological phenomena, and plays crucial roles in epigenetic regulation of eukaryotic gene expression. It is also suggested that DNA methylation alters the mechanical properties of DNA molecules, which is likely to affect epigenetic regulation. However, it has not been systematically investigated how methylation changes the structural and dynamic features of DNA. In this research, to elucidate the effects of methylation on DNA mechanics, a fully atomic molecular dynamics simulation of double-stranded DNA with several methylation patterns was performed. Through the analysis of the relative positioning of the nucleotides (base-step variables), characteristic changes in terms of local flexibility were observed, which further affected the overall DNA geometry and stiffness. These findings may serve as a basis for a discussion on methylation-dependent DNA dynamics in physiological conditions.

## I. INTRODUCTION

Methylation is a universal biochemical modification of DNA [1–3]. A methyl group (− CH_3_) binds to the cytosine base in a CpG dinucleotide, referred to as mCpG [1]. DNA methylation is a widespread and crucial epigenetic modification that influences gene expression [2, 4]. Vertebrate genomes are globally methylated at CpG dinucleotides; only CpG islands (CGI) are unmethylated [5]. CGIs are short unique DNA sequences that are over 200 bp (typically 500–1000 bp) in length and have high G+C content, as well as high CpG density [6]. The promoter regions of 60–70 % of all vertebrate genes overlap with CGIs [7, 8]. CGIs eliminate nucleosomes and attract proteins that create a transcriptionally permissive chromatin state [6, 9]. Whereas, when DNA methylation occurs at CpG sites, the CGI-promoter strongly silences transcription by inhibiting transcriptional factor binding or recruitment of mCpG binding proteins and co-repressor complexes [10]. It has been proposed that methylated CGI functions to stably maintain a silenced state of gene transcription (called the “long-term locking model”) [4].

DNA methylation affects the mechanical properties of DNA. For example, DNA methylation reduces the flexibility of DNA [11], inhibits or facilitates DNA strand separation [12], attracts homologous DNA segments [13], and changes the dynamics of nucleosomes [14]. DNA methylation is comparable to the sequence-dependent mechanics of DNA [15, 16], which affects basal processes of molecular recognition [17]. The extremely opposite roles of CGIs in transcriptional regulation are likely to be defined by the mechanical properties of hyper- and hypo-methylated DNA. However, how DNA methylation changes the structural and dynamic features of DNA molecules remains poorly understood. Although effects of methylation in general on DNA dynamics have been recently studied using molecular dynamics (MD) simulations [18–22], there still remain microscopic properties (e.g. positional effects and their ranges) to be elucidated; thus, systematic analysis of different patterns of methylation is desired.

In this research, the relationship between methylation patterns and DNA dynamics was investigated using all-atom MD simulations. The structures of DNA with several methylation patterns were sampled in equilibrium. Through the analysis of fluctuations in relative positioning of nucleotides, characteristic changes of base-step parameters (BSP) in terms of local flexibility were identified, which further affected overall DNA dynamics.

## II. MATERIALS AND METHODS

### A. Simulation Procedure

All-atom MD simulations were employed for double-stranded DNA dynamics in solution. DNA structures were constructed using X3DNA [23, 24] with a helical parameter set obtained by *in vitro* experiments and X-ray crystal structure analysis [15, 25, 26]. The atomic coordinates of double-stranded DNA were generated according to a given DNA sequence (see [23, 24] for details). In this study, for simplicity, repeats of CpG dinucleotides were used, and regular patterns of methylation were introduced, as shown in Tab. I. These DNA models were soaked in a 60 Å × 60 Å × 200 Å water box, neutralized by K^+^, and added 150 mM KCl. TIP3P water model was employed. VMD [27] was used to infer missing atom coordinates, solvate the model, and visualize the structure throughout the study.

**TABLE I.** Target Sequences. For each model, methylated CpG (mCpG) content ratio, methylated cytosine (mC) sites, and repeat units are listed. All sequences are composed of only CpG dinucleotides. mC sites are indexed as shown in Fig. 1 (c). Note that the models, except SEQ 0.00 and SEQ 1.00, include partial (fractional) sequences at the ends (see Tab. S1 [30]).

| Model | mCpG Content | mC Site (Nucleotide Indices) | Repeat Unit |
| --- | --- | --- | --- |
| SEQ 0.00 | 0% (0) | - | CG |
| SEQ 0.25 | 25% (1/4) | 1 9 17 25 33 41 49 51 59 67 75 83 91 99 | CGCGmCGCG |
| SEQ 0.33 | 33% (1/3) | 1 7 13 19 25 31 37 43 49 51 57 63 69 75 81 87 93 99 | CGmCGCG |
| SEQ 0.50 | 50% (1/2) | 1 5 9 13 17 21 25 29 33 37 41 45 49 51 55 59 63 67 71 75 79 83 87 91 95 99 | CGmCG |
| SEQ 0.67 | 67% (2/3) | 3 5 9 11 15 17 21 23 27 29 33 35 39 41 45 47<br>53 55 59 61 65 67 71 73 77 79 83 85 89 91 95 97 | CGmCGmCG |
| SEQ 0.75 | 75% (3/4) | 3 5 7 11 13 15 19 21 23 27 29 31 35 37 39 43 45 47<br>53 55 57 61 63 65 69 71 73 77 79 81 85 87 89 93 95 97 | CGmCGmCGmCG |
| SEQ 1.00 | 100% (1) | 1 3 5 7 9 11 13 15 17 19 21 23 25 27 29 31 33 35 37 39 41 43 45 47 49<br>51 53 55 57 59 61 63 65 67 69 71 73 75 77 79 81 83 85 87 89 91 93 95 97 99 | mCG |

All simulations were performed using NAMD (version 2.13 multi-core with CUDA) [28, 29]. The CHARMM36 force-field (July 2017 update) was used and a 5MC2 nucleotide patch was applied to introduce 5-methylated cytosine (mC) (Fig. 1 (a)). A periodic boundary condition with Particle-Mesh Ewald electrostatics was employed and a cutoff of 12 Å (with switching from 10 Å) was used for non-bonded interactions. Temperature and pressure were set at 300 K and 1 atm, respectively; a Langevin thermostat (damping coefficient: 5/ps) and Langevin-piston barostat were adopted. After energy minimization (10,000 steps), the system was equilibrated for 10 ns, and then simulated for 50 ns (time-step: 2 fs). The conformation was sampled over the last 50 ns at 10 ps intervals (i.e. 5,000 snapshots), and each DNA model was simulated 10 times (i.e. 50,000 snapshots in total).

**FIG. 1.**
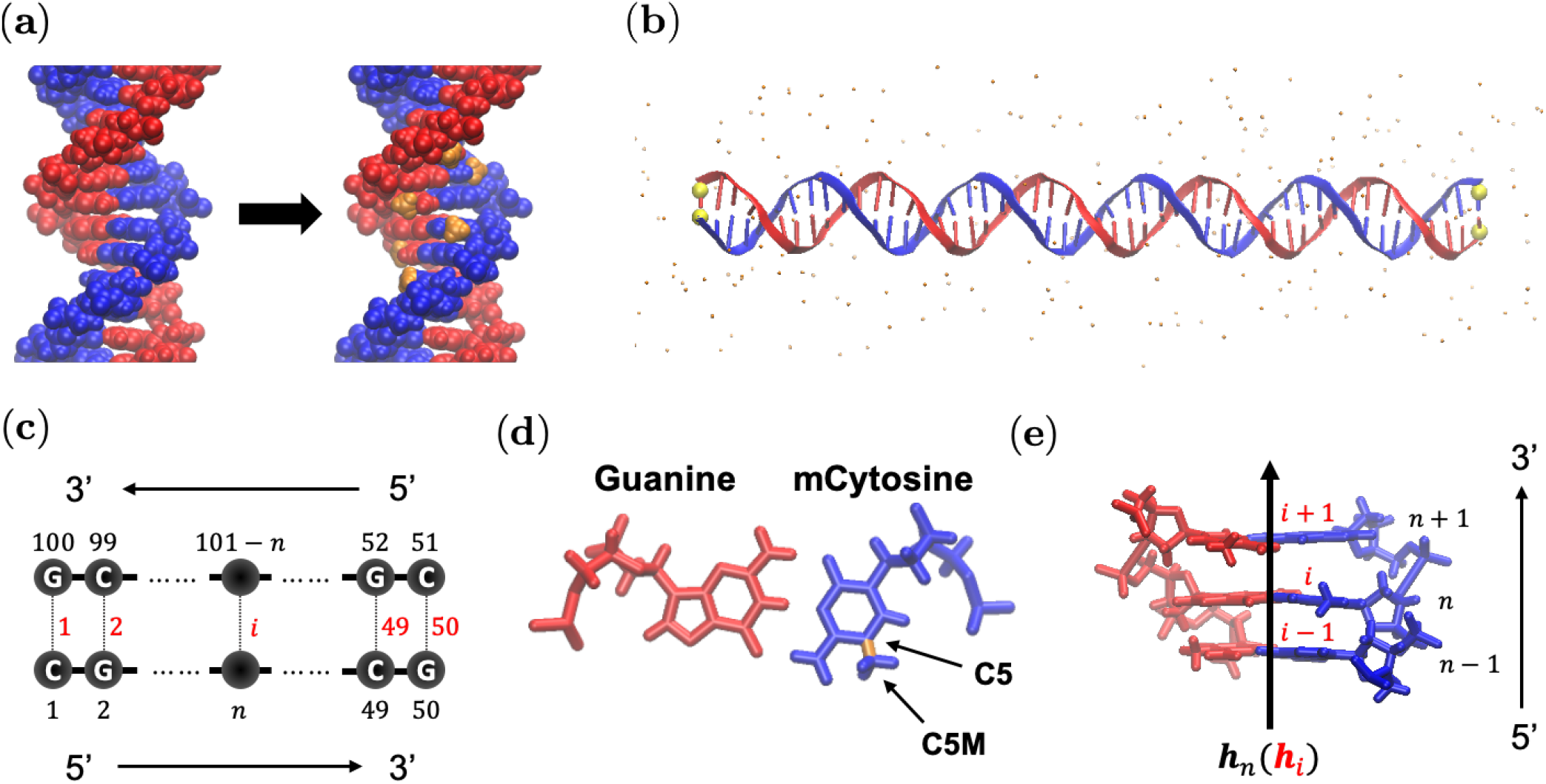
Overview of the Model. (a) Structure of DNA with poly-CpG (left) and poly-mCpG (right). The methyl groups are shown in orange. (b) Solvated DNA model. Small particles represent ions. C1’ atoms located at both ends (shown in yellow) are restrained in the simulation (see Sec. II A). (c) Nucleotide indices *n* (shown in black) and base-pair indices *i* (shown in red) of the double-stranded DNA. (d) Locations of C5 and C5M atoms in 5-methylated cytosine. (e) Schematic representation of helical axis ***h**_n_* (***h**_i_*).

Through the equilibration and production run, harmonic restraints were applied to C1’ atoms of nucleotides at both ends of the DNA segment (Fig. 1 (b); spring constant: 1.0 pN/Å, centered at the position after the energy minimization), to prevent rotation of DNA. It was confirmed that the restraints did not excessively stretch the DNA and allowed bending (Fig. 2 (c)).

**FIG. 2.**
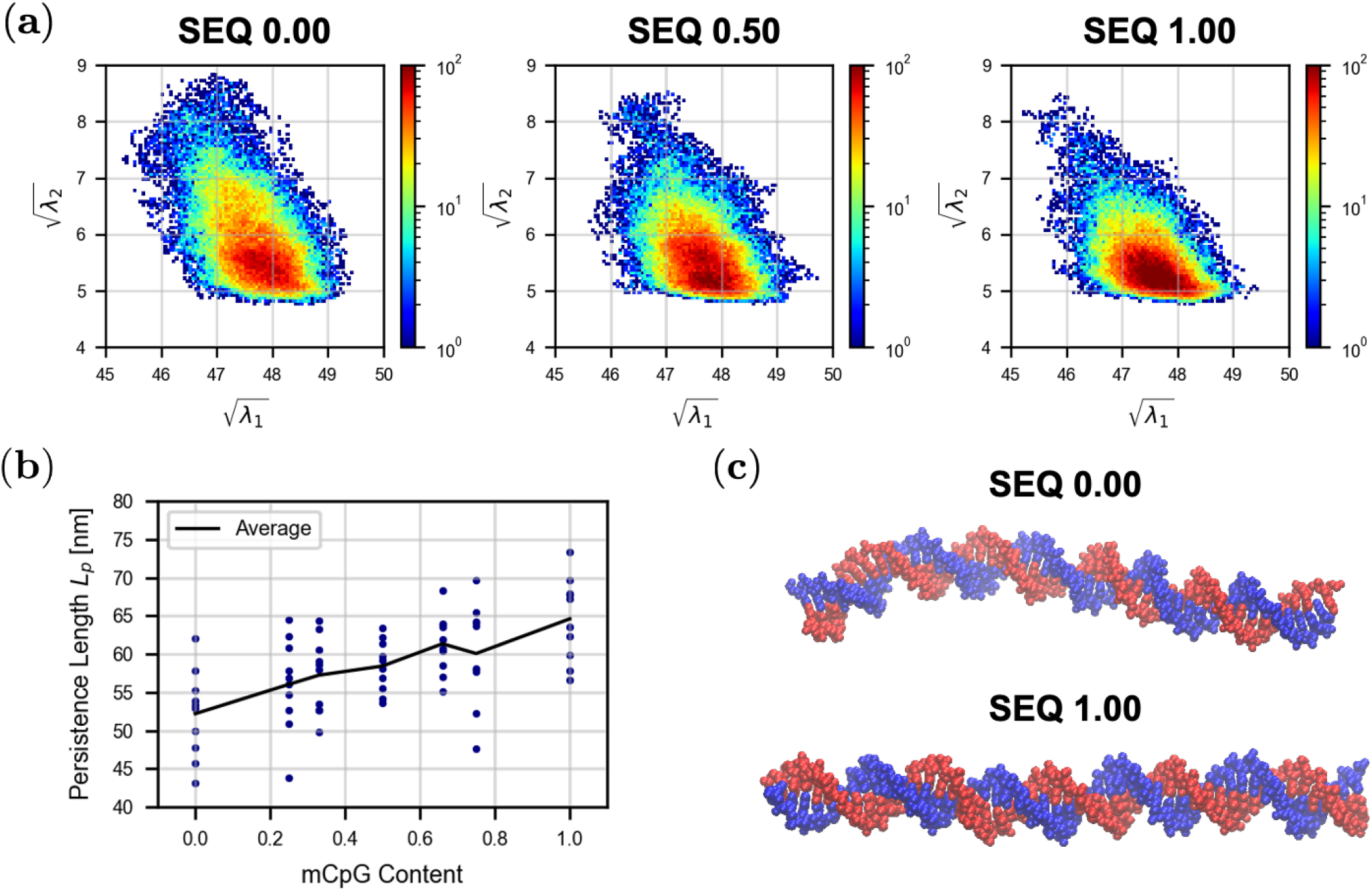
Profiles of Overall Geometry and the Persistence Length of DNA. (a) Overall geometry. The distribution of (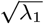, 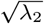) summed over 10 trials is shown. (b) Persistence length *L_p_*. The average within each trial and over 10 trials are shown. The horizontal axis corresponds to the mCpG content (Tab. I). *p*-values (by Welch’s *t*-test) between mCpG content 0.0 and 0.5, 0.5 and 1.0, and 0.0 and 1.0 are 9 × 10^−3^, 8 × 10^−3^, and 1 × 10^−4^, respectively. (c) Snapshots of the structure of SEQ 0.00 and SEQ 1.00 models at 40 ns.

### B. Overall Geometry of Double-Stranded DNA

Overall structural variation of the DNA model was characterized by the ratios of the square root of the three principal components of the distribution of atomic positions (except hydrogen), 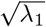, 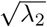, and 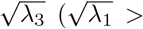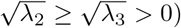. Here, *λ*_1_ to *λ*_3_ were obtained as eigen-values of the covariance matrix *I*:

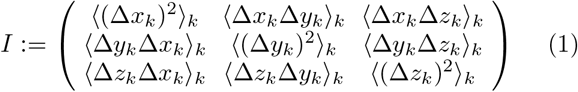

where (Δ*x_k_*, Δ*y_k_*, Δ*z_k_*) = (*x_k_* − ⟨*x_m_⟩_m_*, ⟨*y_k_* − ⟨*y_m_⟩_m_*, *z_k_* − ⟨*z_m_⟩_m_*); (*x_k_, y_k_, z_k_*) is the position of *k*-th atom, (⟨*x_m_⟩_m_*, ⟨*y_m_⟩_m_*, ⟨*z_m_⟩_m_*) is the center position of a given DNA molecule, and ⟨…⟩*_m_* indicates the average for all *m*s. Note that (*λ*_1_, *λ*_2_, *λ*_3_) were defined for each snapshot *τ*, and thus 5,000 data-points were obtained in each simulation trajectory (see Sec. II A).

In this case, *λ*_1_ and *λ*_2_ corresponded to the extension and distortion of double-stranded DNA, respectively.

### C. Persistence Length of Double-Stranded DNA

The persistence length of polymers statistically characterize their stiffness [31], and the case of double-stranded DNA has been studied well [32–34]. In this research, the persistence length of DNA was defined as follows. First, *A_d_*, the inner-product of helical axes separated by *d* nm (*d* ∈ [0.3, 10.0] at intervals Δ*d* (= 0.01)) was defined as

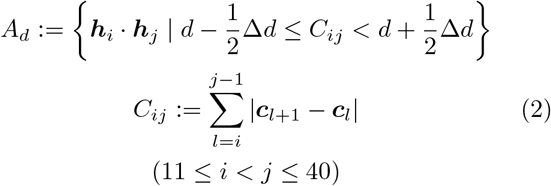

where ***c**_i_* is the center of base pair *i*, and ***h**_i_* is the helical axis of base-pair *i* (Fig. 1 (e)). These vectors were determined using X3DNA [23, 24]. The length of ***h**_i_* was equal to 1. To eliminate the effects of the boundaries and restraints (see Sec. II A), 10 base pairs from each end were eliminated (the base-pair indices (*i, j*) satisfy 11 ≤ *i* < *j* ≤ 40). Then, the dataset 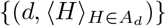 (⟨…⟩_*H*_, which indicates the average for all *H*s, sampled over all 5,000 snapshots for each trial) was fitted by function *f*(*d*) := exp(− *d/L_p_*), where *L_p_* is the persistence length to be estimated.

### D. Local Geometry and Flexibility

Profiles of base-step geometry were obtained using X3DNA [23, 24], from which the local flexibility was estimated. First, the displacement in terms of mode represented by each base-step parameter (BSP) from the (meta)stable conformation was examined. The score 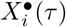 for BSP • of base-step *i* (base pair indices from *i* to *i* + 1 shown in Fig. 1 (c)) at the *τ*-th frame, and its displacement 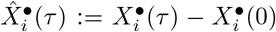 were calculated, where 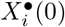 is the score 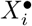 just after the energy minimization (*τ* = 0) (see Sec. II A). Then, the flexibility score 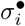 for BSP • of base-step *i* was defined as the standard deviation (S.D.) of 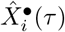, either in each simulation trial or over 10 trials for the same model.

Additionally, to quantify the positional effects of methylation (i.e. effects depending on the relative position to the mCpG sites) on the structural variation, the flexibility scores 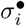 were aligned and averaged in terms of the repeat unit (Tab. I), to obtain 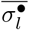 defined in Eq. (3). The repeat units and partial (fractional) sequences at the ends for each model are shown in Tab. S1 [30]. To eliminate boundary effects, 10 base-pairs were omitted from each end, and only complete repeat units within the central 30 base-pairs were analyzed (for base-step parameters, the base-step between the last base-pair to the next one was also included). For example, for SEQ 0.33, five iterations of CGmCGCG in the center (and the next C for base-step parameters) were used. The averaged scores 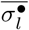 at base-step *l* in the repeat unit were calculated as

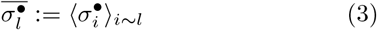

where ⟨…⟩_*i*~*l*_ represents the average for all base-step *i*s that correspond to position *l* in the repeat unit (Tab. I). 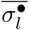 were calculated for each trial, and then their mean and S.D. over 10 trials were evaluated, for which Welch’s *t*-test was applied.

Furthermore, relationships between 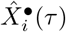 were evaluated. Specifically, (i) the same BSP • between two base-steps, and (ii) two BSPs • of the same base-step, were compared. For case (i), correlation coefficient of BSP • between base-steps *i* and *j* was calculated as

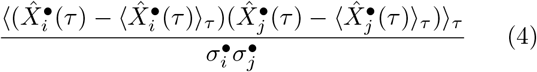

where ⟨…⟩_*τ*_ indicates the average for all *τ*s over 10 trials (50,000 samples in total, for which 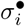 and 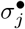 were also calculated) for each model. For case (ii), correlation coefficient between two individual BSPs •_1_ and •_2_ (of base-step *i*) was calculated as

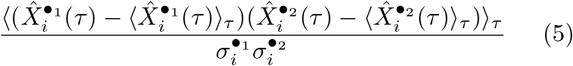

where ⟨· · ·⟩_*τ*_ is the same as in Eq. (4). Then, it was averaged over the central 30 base-steps (11 ≤ *i* ≤ 40).

### E. Dynamics of Methyl Groups

The relative orientation of each methyl group was evaluated by the angle between the direction of the methyl group and the plane formed by the base-pair. The angle *θ_n_* was defined as follows:

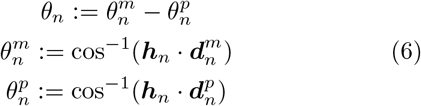

where ***h**_n_* is the vector of the helical axis at the base-pair where nucleotide *n* belongs (Fig. 1 (e)), obtained using X3DNA [23, 24], 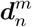 is the direction vector of the methyl group (from C5 to C5M shown in Fig. 1 (d)), and 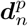 is the direction vector from the C1’ atom to the C1’ atom in the complementary base. The lengths of these three vectors were equal to 1. If *θ_n_* > 0 (*θ_n_* < 0), the methyl group tilted toward the next (prior) base.

Additionally, physical contact (interaction) frequency between the methyl group and its neighbor nucleotides was evaluated. A methyl group and a nucleotide were regarded in contact if their shortest atomic distance was less than 3.0 Å.

## III. RESULTS

### A. Profiles of Double-Stranded DNA Stiffness

The over all geometry of DNA was represented by the extension 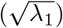 and distortion 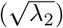 of its atomic coordinates. The distribution of 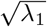 and 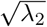 for each model is shown in Figs. 2 (a) and S1 [30]. Mostly, (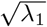, 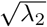) was in the range of 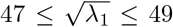 and 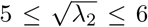, corresponding to a conformation similar to the shape of stretched DNA (e.g. Fig. 1 (b)).

The distribution outside of this range was different for these models. The probability within this range was higher for SEQ 1.00 than that for SEQ 0.00. 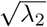 of SEQ 1.00 tended to be smaller than that of SEQ 0.00. This implied that high mCpG content resulted in a stretched, or stiffer DNA shape. The other models (Tab. I) showed intermediate stiffness between SEQ 0.00 and SEQ 1.00 (Figs. 2 (a) and S1 [30]). The variation of 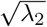 was suppressed as the mCpG content increased. These results showed that higher methylation content stiffened DNA in equilibrium conditions.

The persistence length *L_p_* (see Sec. II C) for each model is shown in Fig. 2 (b), which increases approximately linearly with mCpG content. Though estimated *L_p_* values showed large variation, this trend was significant (e.g., *p* < 1 × 10^−4^ between SEQ 0.00 and SEQ 1.00). Note that the typical persistence length of DNA is approximately 50 nm (~ 150 base-pairs), with which the estimated values are consistent.

The difference in DNA stiffness can also be observed in the snapshots, as shown in Fig. 2 (c).

### B. Local Base-Step Flexibility

The overall DNA geometry analyzed above is a result of sequential base-pair stacking [35, 36]. To investigate the effects of methylation in detail, local profiles of flexibility at each base-step were evaluated. Through the analysis using X3DNA (see Sec. II D), profiles of BSPs were obtained (Fig. S2 [30]). Profiles of 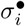 (Fig. S2 [30]) changed with increase of mCpG content; among them, shift, tilt, and twist (shown in Fig. 3) were the most significantly affected.

**FIG. 3.**
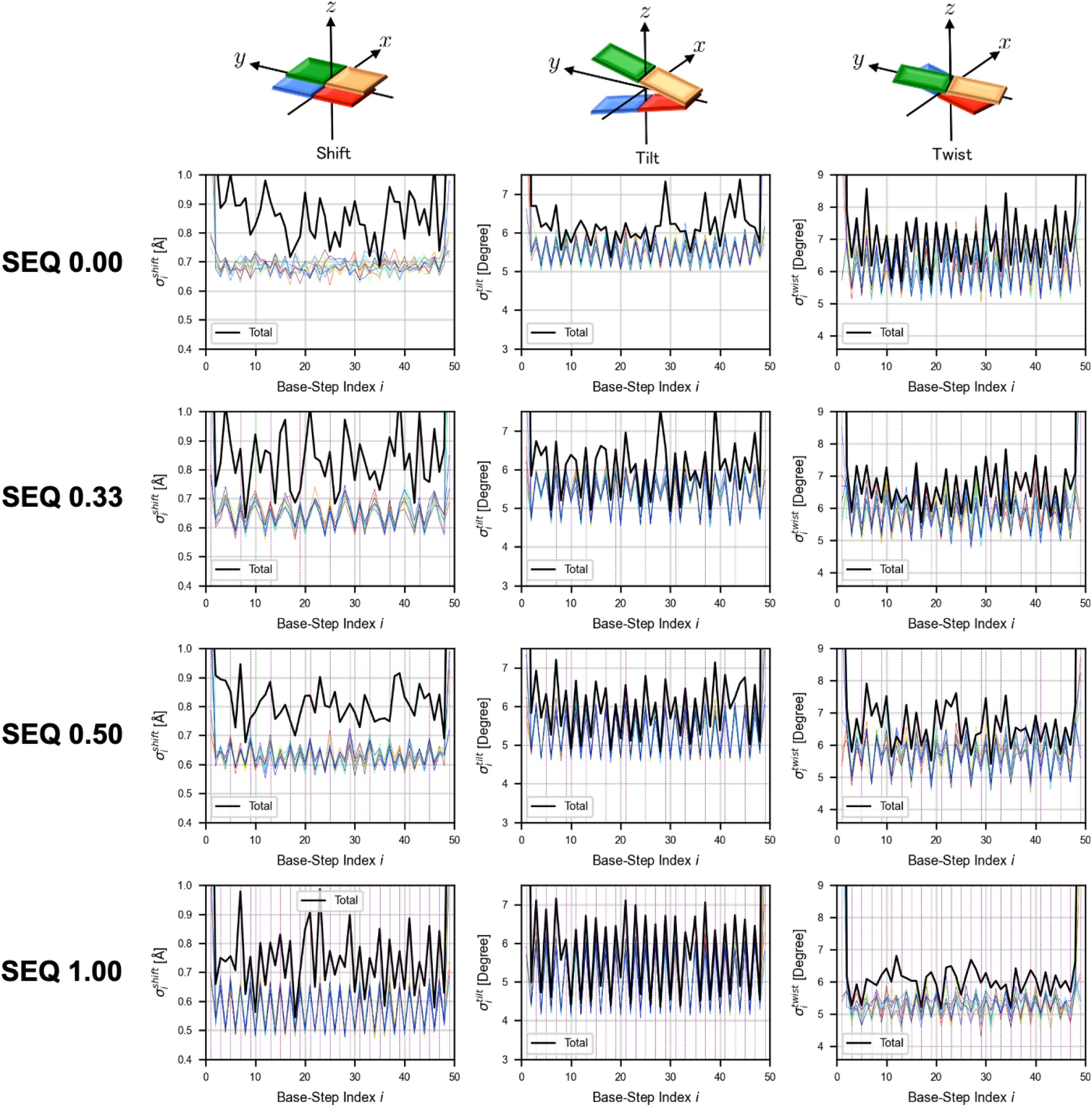
Profiles of 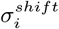, 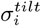, and 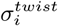. Colors show different simulation trials and the black line shows the overall flexibility (structural variation) in 10 trials. In the horizontal axis, odd and even numbers correspond to base steps of C→G and G→C, respectively (see Fig. 1 (c)). Purple lines show mC→G (C→G in the case of mC) base steps.

In the profiles of 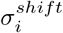 and 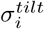 shown in Fig. 3, colored curves (i.e. 10 individual simulation trials) show alternating patterns, getting steeper as mCpG content increases. More specifically, with higher mCpG content, 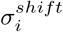 and 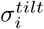 at base-steps G→C became smaller, while that at C → G was almost unchanged. The overall 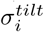 for 10 trials clearly showed alternating patterns similar to individual trials as mCpG content increased, while 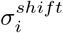 did not show such clear patterns. Taking into account the results of partially methylated cases (e.g. SEQ 0.50), methylated DNA made the tilt-mode stiffer at base-steps G → C, and restricted their deformation in determined ranges (Fig. 3).

Furthermore, mCpG suppressed tilt and shift dynamics only at the base-steps adjacent to the mCpG. To clearly show the positional effect, the profiles of 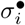 in terms of the repeat unit (i.e. relative position with respect to the mCpG) have been aligned and averaged (see definition of 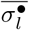 in Sec. II D), as shown in Figs. 4 and S3 [30]. In the profile of SEQ 0.33 in Fig. 4 (a), 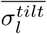 decreases at both neighbors of mCpG (Tab. I), while almost unchanged at G→C between non-methylated CpGs (indicated by ↓^×^ in Fig. 4).

**FIG. 4.**
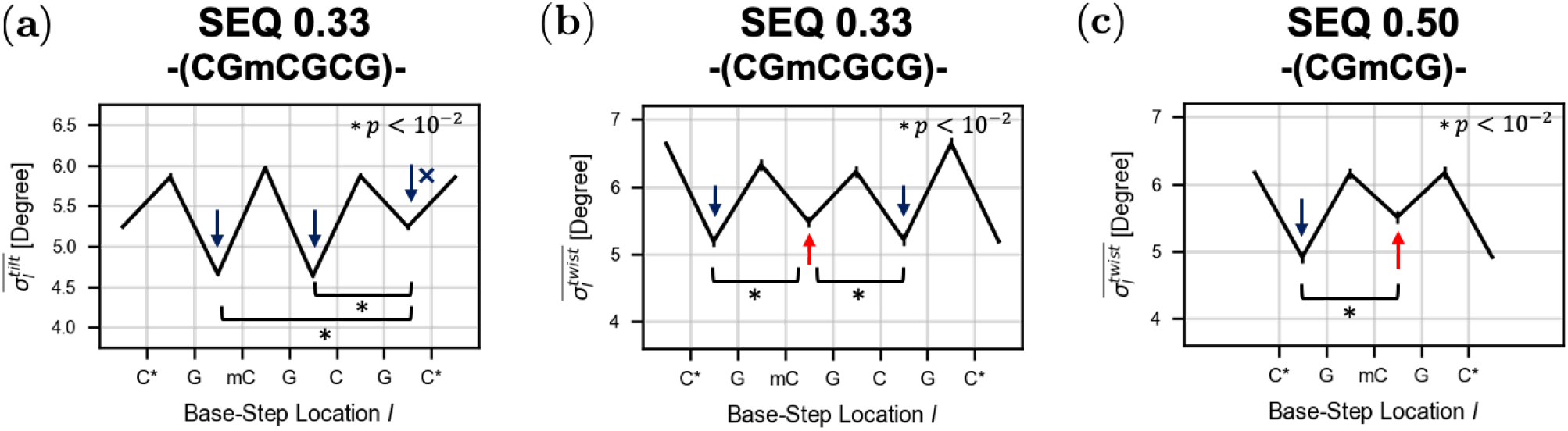
Profiles of 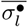 for the Repeat Unit. 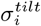 and 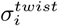 (shown in Fig. 3) are aligned and averaged in terms of the repeat unit (Tab. I). Error bars show mean ± S.D. over 10 trials; see Sec. II D for the method. As these profiles are for base-step parameters, data points are shown between two consecutive bases. Nucleotides at both ends (shown with * in the horizontal axis) are identical. (a) 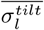 of SEQ 0.33 (CGmCGCG, averaged over five iterations); (b) 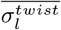 of SEQ 0.33 (CGmCGCG, *l* averaged over five iterations); (c) 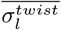 of SEQ 0.50 (CGmCG, averaged over seven iterations). In (a), reduction of S.D. at G→C is significantly larger at both neighbors of mCpG (* *p* = 3 × 10^−16^ each). In (b) and (c), S.D. at mC→G is significantly higher than unmethylated C→G (* *p* = 1 × 10^−7^ (b; left), 1 × 10^−6^ (b; right), and 6 × 10^−11^ (c)).

In contrast, for 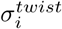, alternating patterns became less significant as mCpG content increased (Fig. 3). In SEQ 0.00, 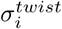 at base-steps G→C was larger than that at C → G, both in individual trials and the overall profile. In SEQ 1.00, the difference between G → C and C → G was smaller, and almost disappeared in the overall 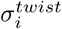 profile. This result suggested that DNA methylation irregularly prevented twist dynamics. In general, DNA methylation made twisting at base-steps G → C stiffer; however, another considerable trend existed. In SEQ 0.50, 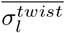 was actually higher at mCpG sites than at non-methylated CpG sites (Fig. 4 (c); indicated by red arrow); this trend also appeared in SEQ 0.33, although it was slight (Fig. 4 (b)). This result suggested that, in terms of twisting, methylation generally stiffened DNA; however, C → G in the mCpG itself showed relatively large fluctuations.

Besides the positional effects on each BSP, correlations between BSPs were surveyed (see Sec. II D), to detect long-range interactions or characteristic motion. However, notable results were not found (Figs. S4 and S5 [30]). Correlation of the same BSP between two base-steps (Fig. S4) was high only for the (near-)diagonal components (i.e. nearby base-steps). Some correlation between BSPs within the same base-step (Fig. S5) was observed, reflecting the structure of DNA (e.g. between roll and twist), which was however independent of methylation.

### C. Methyl Groups Distribution

The orientation of methyl group *θ_n_* (see Sec. II E) was evaluated (Figs. 5 (a) and S6 [30]). In every case, the *θ_n_* distribution was both positive and negative, and the aver-age was approximately −0.2 [radian]. This suggested that methyl groups slightly tilt toward the 5’-end, but could fluctuate. No specific interactions (e.g. electrostatic interactions) were identified between the methyl group and other parts of the DNA molecule; instead, physical contacts between methyl groups and their 5’-side neighbor nucleotides were frequently observed (Figs. 5 (b) and S7 [30]). Hence, it was suggested that weak van der Waals interactions and simple excluded volume effects played a role in changing local base-step flexibility.

**FIG. 5.**
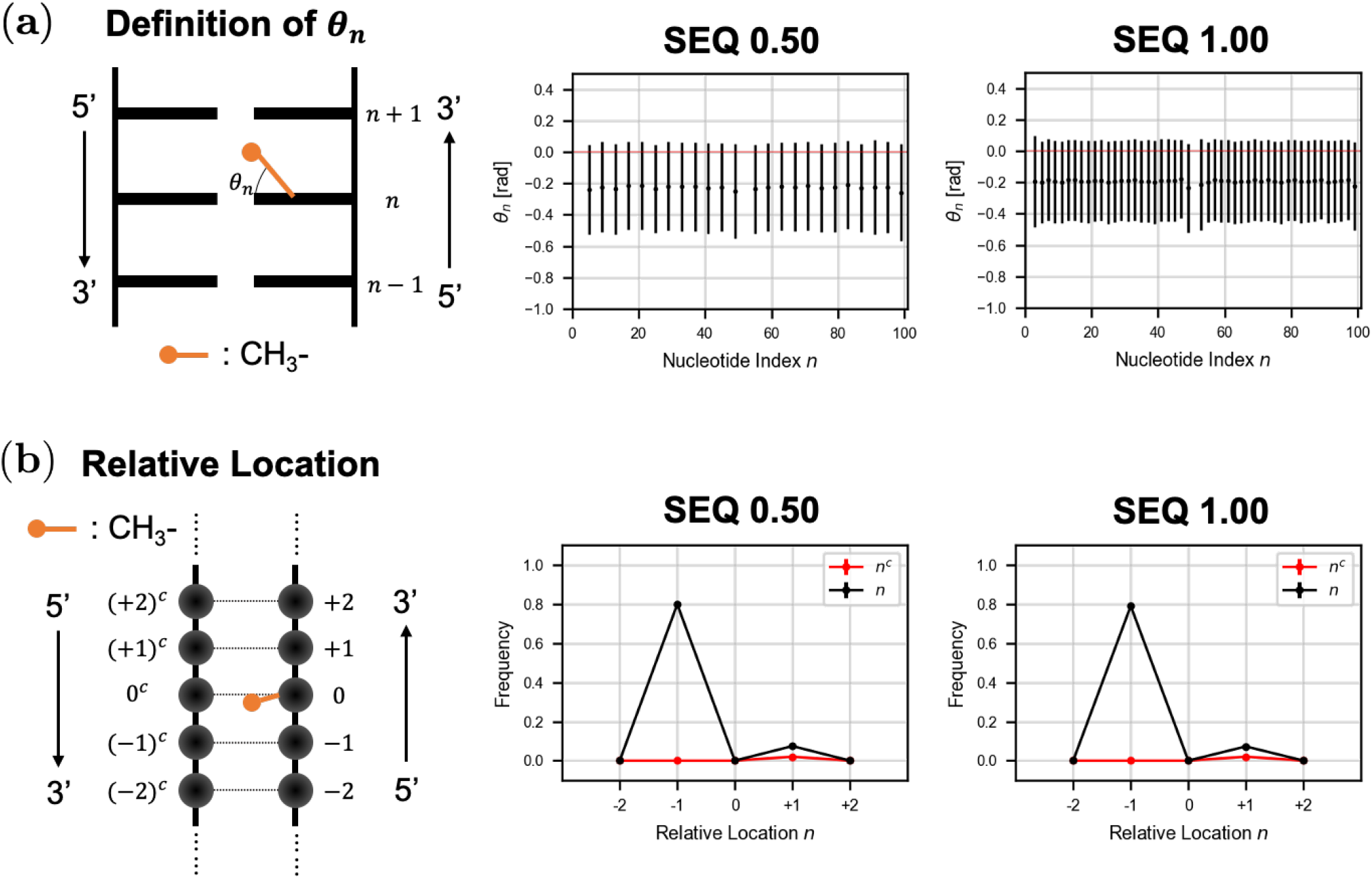
Orientation of Methyl Groups and Their Interactions with DNA. (a) The definition and observed distribution of *θ_n_*. The average ± the standard deviation (S.D.) of *θ_n_* for each methyl group is shown. (b) The definition of nucleotide indices relative to the methyl group, and estimated contact frequencies. The average ± S.D. of contact frequency for all methyl groups is shown.

## IV. DISCUSSION AND CONCLUSIONS

In this study, how different patterns of methylation affected DNA dynamics, both overall geometry (Fig. 2) and local flexibility at each base-step in terms of shift, tilt, or twist (Fig. 3) were investigated. Methylation generally stiffens DNA, which is consistent with previous studies. Specifically, the shifting and tilting dynamics of base-steps adjacent to the mCpG was clearly restricted. Nevertheless, in terms of twisting, C→G base-steps in mCpG were certainly more flexible than other sites. This result might originate from the orientation of the methyl groups and their interactions with neighbors (Fig. 6). The twisting flexibility of C→G, maintained at the mCpG itself, may enhance accessibility of mCpG binding proteins [37, 38], although further investigation is required.

**FIG. 6.**
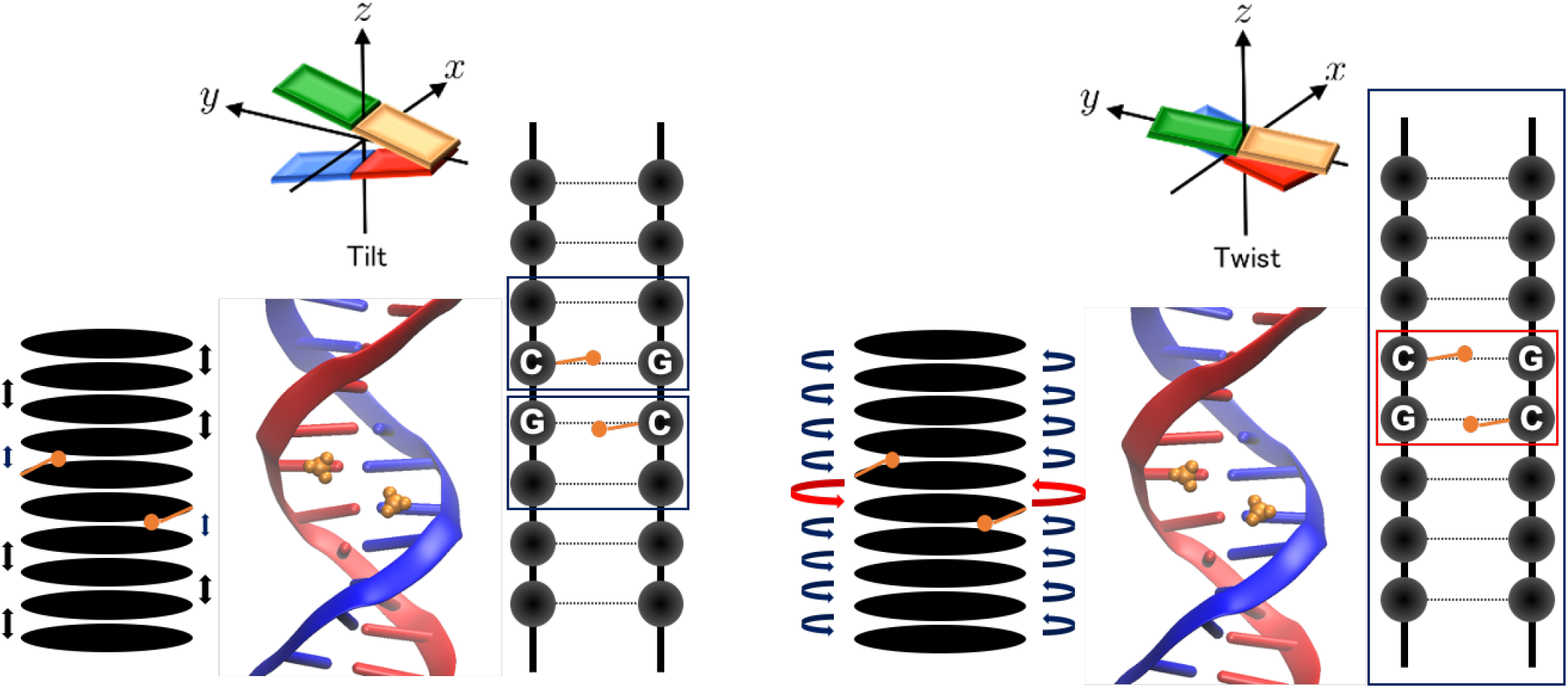
Schematics of Directed Effects Induced by DNA Methylation. Suggested effects on tilt and twist dynamics. Navy (or red) squares represent ranges where site-specific DNA methylation (i.e. existence of mCpG) makes the tilt or twist mode stiffer (or relatively flexible).

Recently, mechanical response of hypermethylated CGI DNA to stretching was assessed by nanometry using optical tweezers [39]; stiffening of DNA was observed, which however disappeared when DNA was over-stretched. In our analysis, it was suggested that the base-step flexibility (Fig. 3) was suppressed by dynamic interactions between DNA and methyl groups (Fig. 6). These interactions are expected only in B-DNA conformation and likely to disappear in the overstretched DNA; hence the suggested mechanism (Fig. 6) is consistent with the experimental result [39]. Although our study suggested the importance of methyl group dynamics (Fig. 5), interactions between DNA and water molecules have not been discussed. Another preceding research suggested that hydration around methylation sites affects local base-step flexibility [22]. Both physical and chemical effects of methylation might be involved in the mechanism.

Previous findings suggest that DNA methylation stabilizes nucleosome positioning [40]. X-ray crystal structures of the nucleosome have been obtained using special sequences of DNA showing high affinity to core histones; in these structures, DNA is not uniformly curved around histones, but bent at several sites [41]. These results also support the hypothesis that site-specific DNA methylation may change local base-step flexibility, which may induce a similar situation and enhance nucleosome positioning.

In this analysis, DNA methylation globally stiffens DNA (Fig. 2), which is consistent with previous experiments [11]. These findings suggest that effects of site-specific methylation on DNA dynamics should be evaluated both at the methylated site and also as global correlated dynamics (e.g. normal modes), as we previously proposed [42, 43]. In recent years, coarse-grained (reduced degree of freedom) physical models of DNA have been proposed, which also include sequence-dependent mechanical properties (e.g. [42–45]). To reproduce the dynamics of longer DNA in physiological conditions, methylation-dependent properties should be also incorporated into these models; this will be performed in future studies.

## DATA AVAILABILITY STATEMENT

The datasets generated for this study can be found in the Zenodo repository DOI: 10.5281/zenodo.3992685.

## AUTHOR CONTRIBUTIONS

TK, MS, AA, and YT conceived and designed the research and wrote the paper. TK, AA, and YT analyzed the data. TK and YT performed the simulations.

## ACKNOWLEDGMENTS

The authors are grateful to I. Nikaido, S. Shinkai, and R. Erban for fruitful discussions. This work was supported by RIKEN Junior Research Associate Program, “TSUBAME Encouragement Program for Young/Female Users” of Global Scientific Information and the Computing Center at Tokyo Institute of Technology, JSPS KAKENHI Grant Numbers JP16H01408, JP18H04720, and 18KK0388, and by JSPS and NRF under the Japan-Korea Basic Scientific Cooperation Program. Part of the numerical calculations were carried out on the TSUBAME 3.0 supercomputer at Tokyo Institute of Technology, and the RIKEN supercomputer HOKUSAI.

## Notes

https://doi.org/10.5281/zenodo.3992685

### Competing Interest Statement

The authors have declared no competing interest.

### Summary of Updates

Figures S4, S5, and S7 added; Figures 2 and 5 revised; Some corrections and additional results in Methods and Results.

