## Supplemental Material for "Structural Dynamics of DNA Depending on Methylation Pattern"

Table S1: Sequence Constitution of the Models. $\star^m$ represents $m$-times of the iteration of the repeat unit.

| Model | Repeat Unit | Sequence Constitution |
|---|---|---|
| SEQ 0.00 | CG | 5'-$\star^{25}$-3' |
| SEQ 0.25 | CGCGmCGCG | 5'-mCGCG-$\star^{5}$-CGCGmCG-3' |
| SEQ 0.33 | CGmCGCG | 5'-mCGCG-$\star^{7}$-CGmCG-3' |
| SEQ 0.50 | CGmCG | 5'-mCG-$\star^{12}$-3' |
| SEQ 0.67 | CGmCGmCG | 5'-$\star^{8}$-CG-3' |
| SEQ 0.75 | CGmCGmCGmCG | 5'-$\star^{6}$-CG-3' |
| SEQ 1.00 | mCG | 5'-$\star^{25}$-3' |

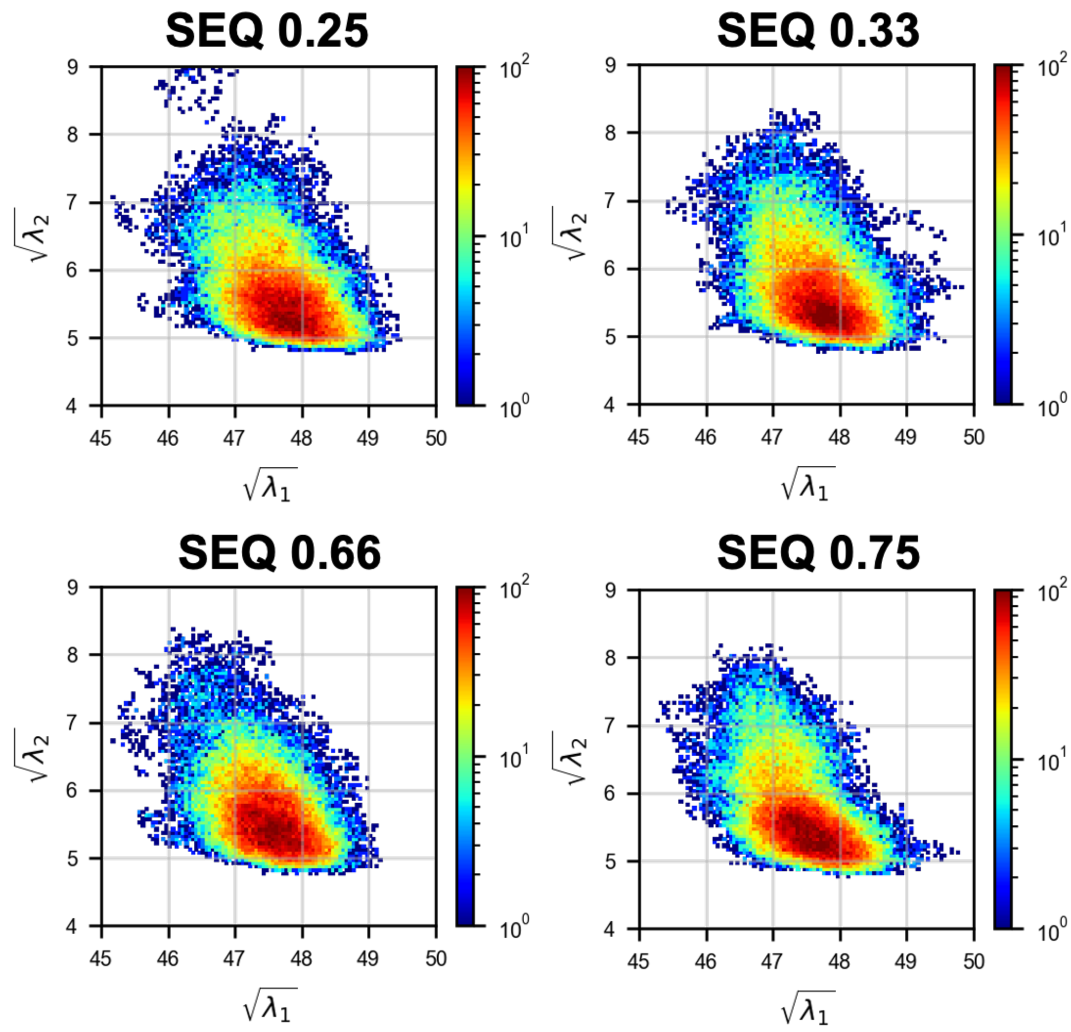


Figure S1: Profiles of Overall Geometry of DNA. The distribution of $(\sqrt{\lambda_1}, \sqrt{\lambda_2})$ summed over 10 trials is shown.

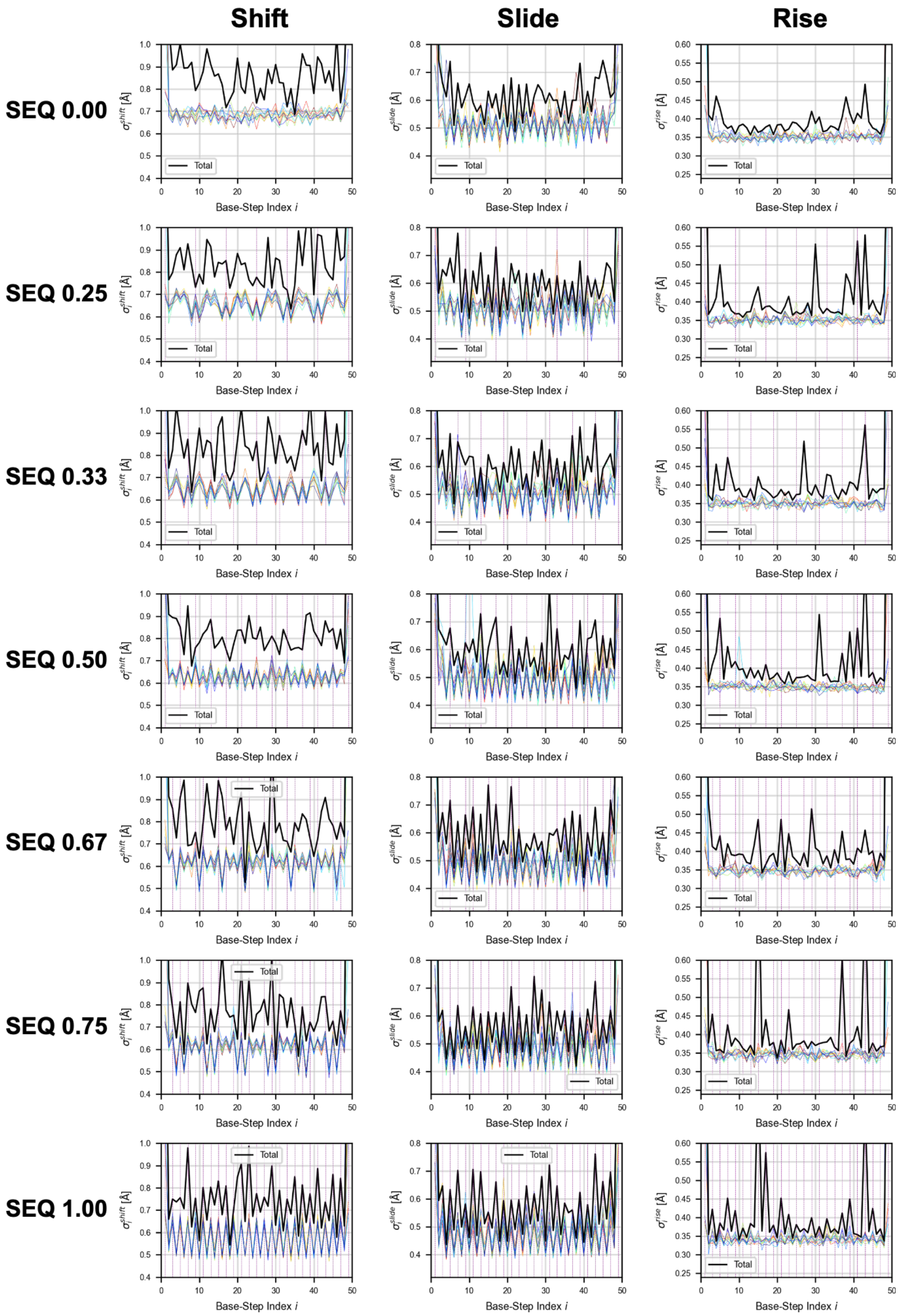


Figure S2: Profiles of $\sigma_i^{\bullet}$. Colors show different simulation trials and the black line shows the overall flexibility (structural variation) in 10 trials. In the horizontal axis, odd and even numbers correspond to base steps of C→G and G→C, respectively (Fig. 1 (c)). Purple lines show mC→G (C→G in the case of mC) base steps.

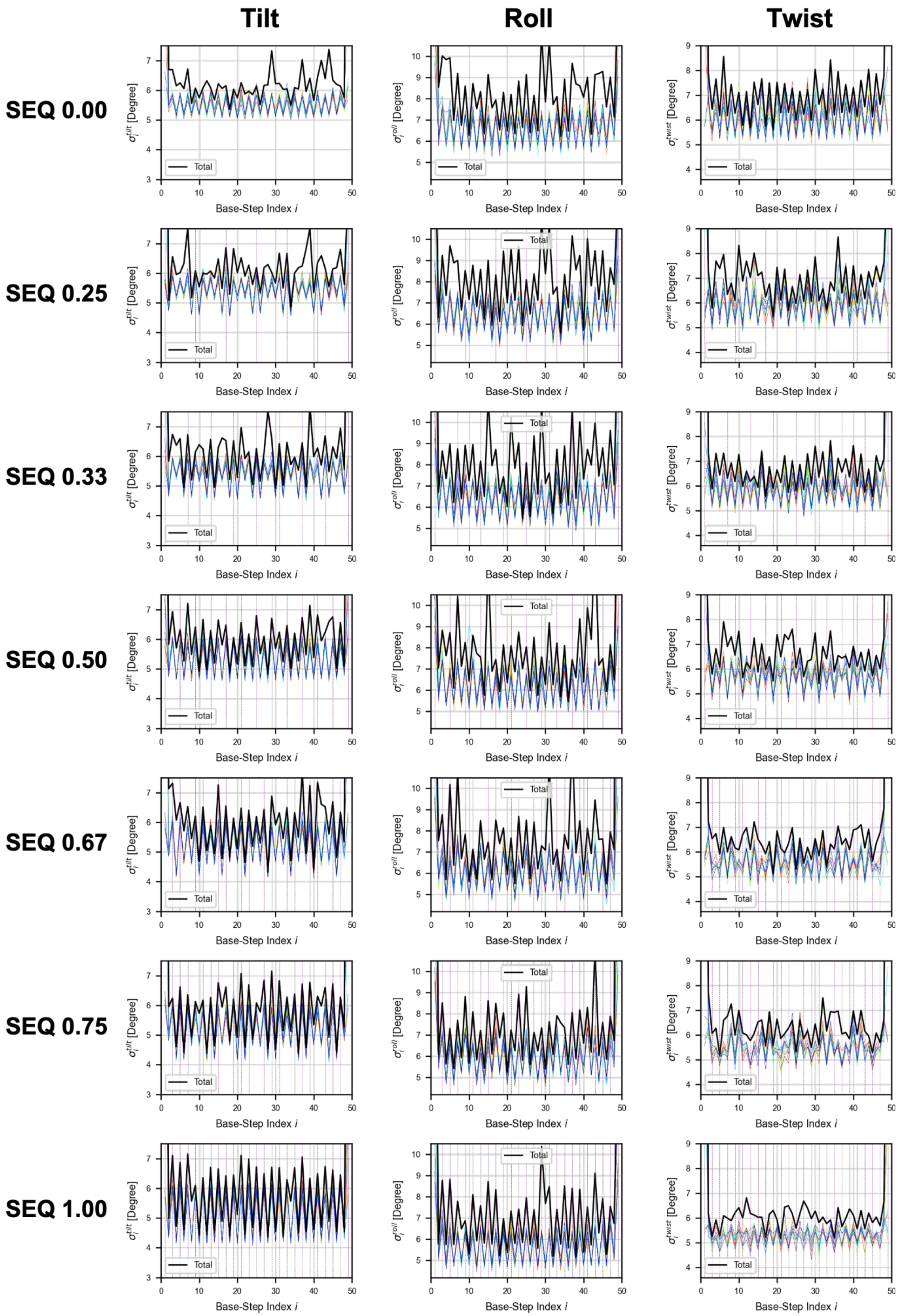
Tilt
Roll
Twist
SEQ 0.00
SEQ 0.25
SEQ 0.33
SEQ 0.50
SEQ 0.67
SEQ 0.75
SEQ 1.00
$\sigma_i^{tilt}$ [Degree]
$\sigma_i^{roll}$ [Degree]
$\sigma_i^{twist}$ [Degree]
Total
Base-Step Index $i$

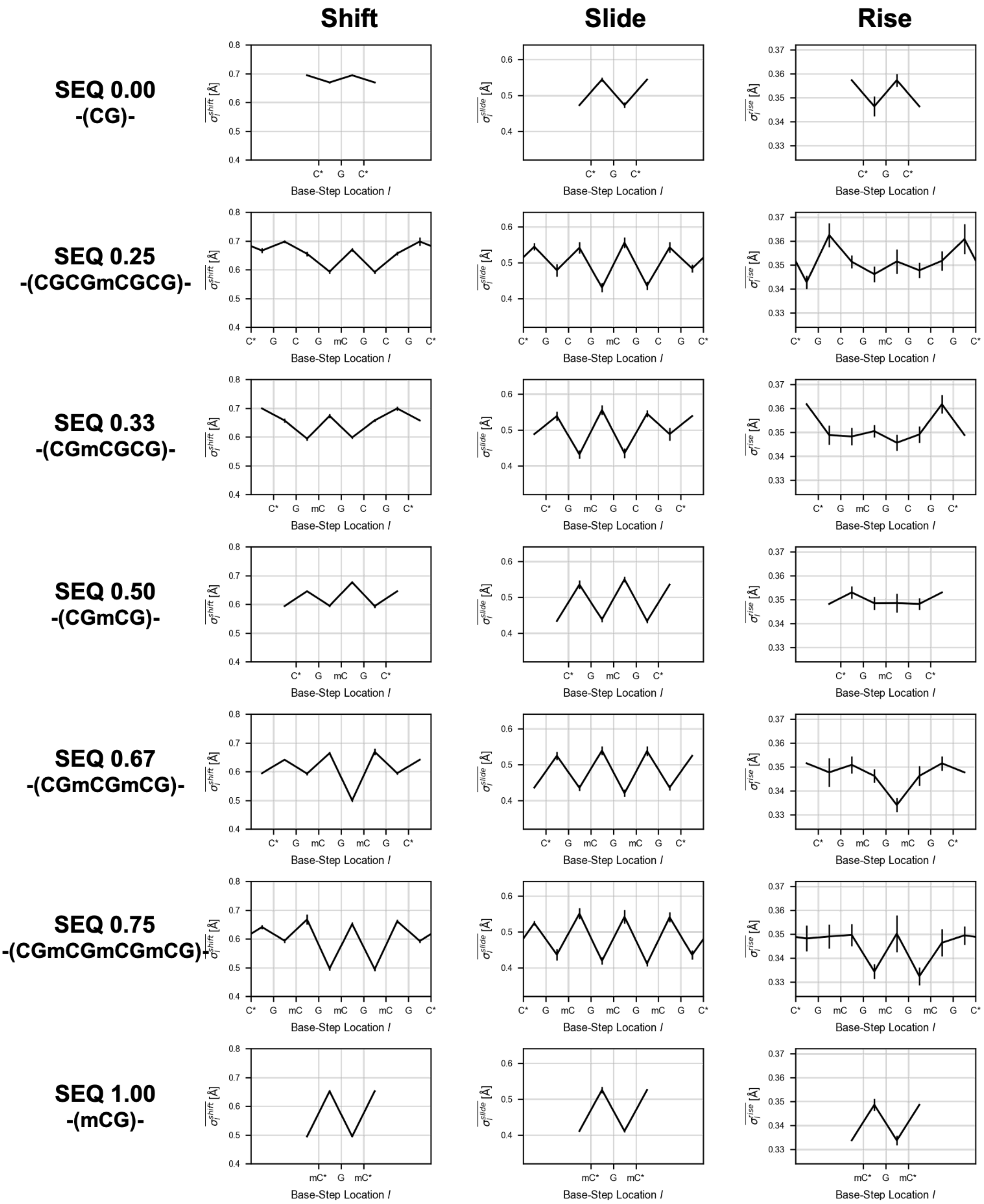


Figure S3: Profiles of $\overline{\sigma_l^\bullet}$ for the Repeat Unit. $\sigma_i^\bullet$ (Fig. S2) are aligned and averaged in terms of the repeat unit (Tab. 1); shown in the same way as in Fig. 4. Nucleotides at both ends (shown with * in the horizontal axis) are identical.

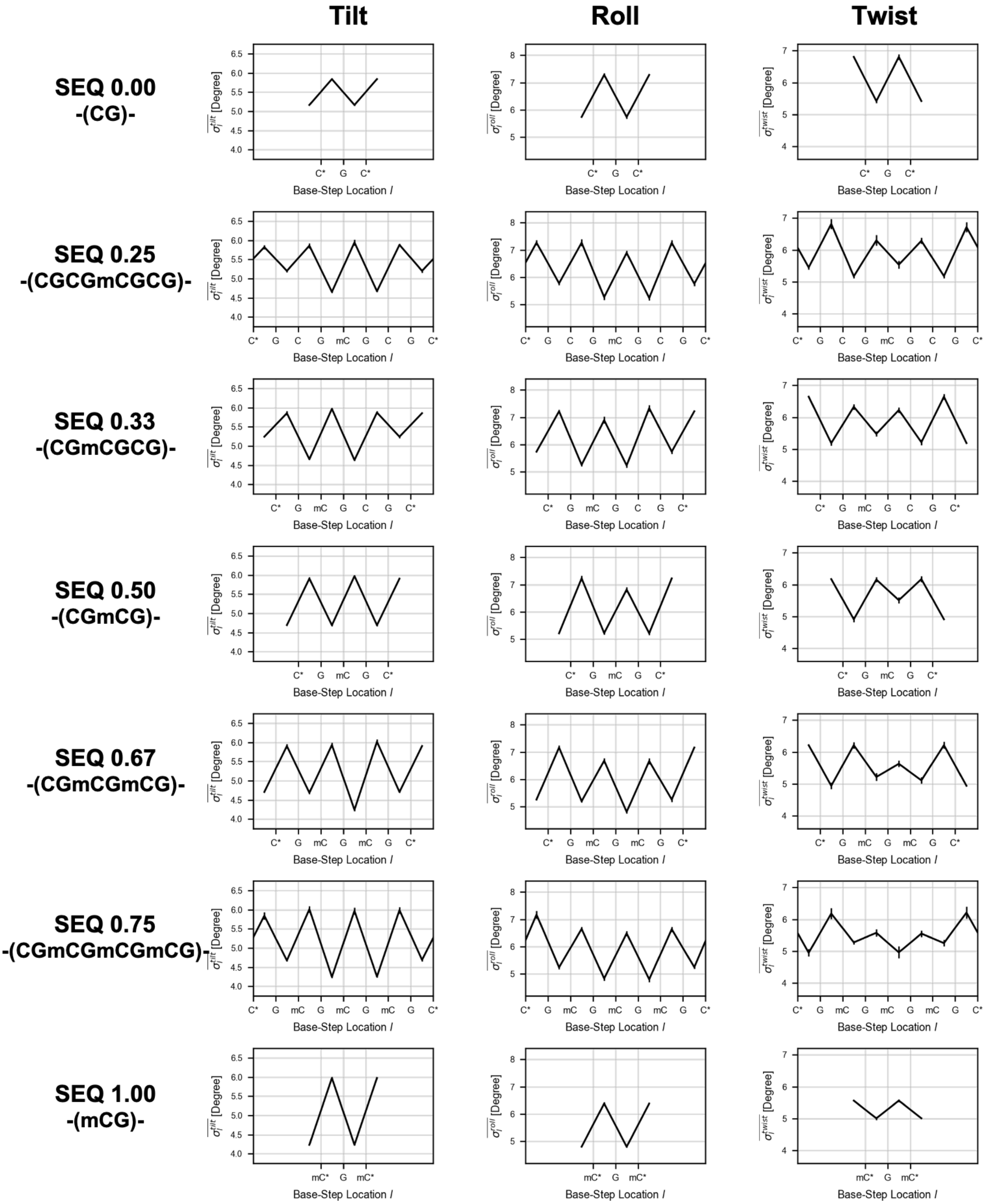
Tilt
Roll
Twist
SEQ 0.00
-(CG)-
SEQ 0.25
-(CGCGmCGCG)-
SEQ 0.33
-(CGmCGCG)-
SEQ 0.50
-(CGmCG)-
SEQ 0.67
-(CGmCGmCG)-
SEQ 0.75
-(CGmCGmCGmCG)-
SEQ 1.00
-(mCG)-
Base-Step Location l
[Degree]

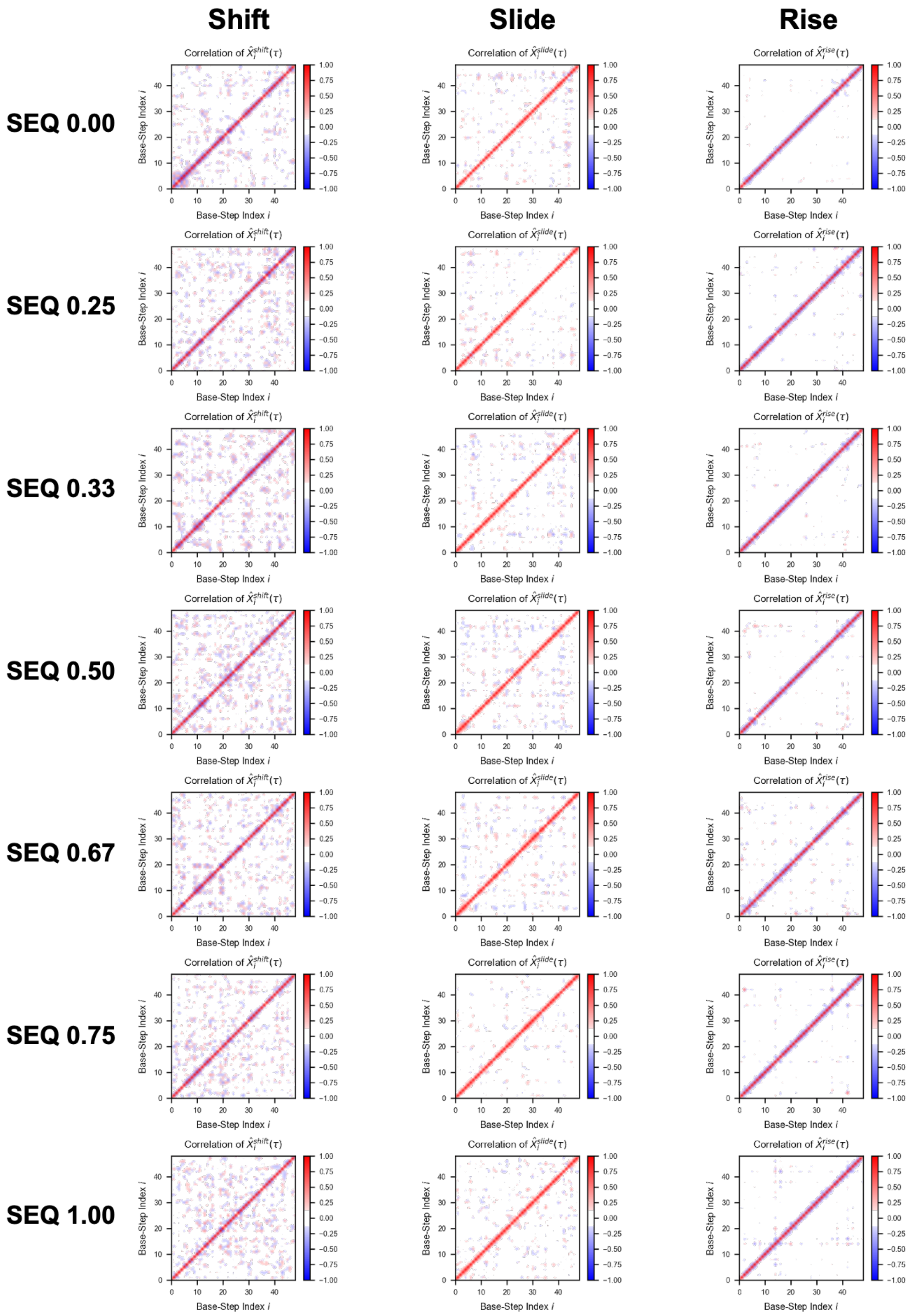


Figure S4: Correlation Coefficient Matrix of $\hat{X}_i^{\bullet}(\tau)$ for Base-step Pairs $(i, j)$. Each matrix component is defined in Sec. II D.

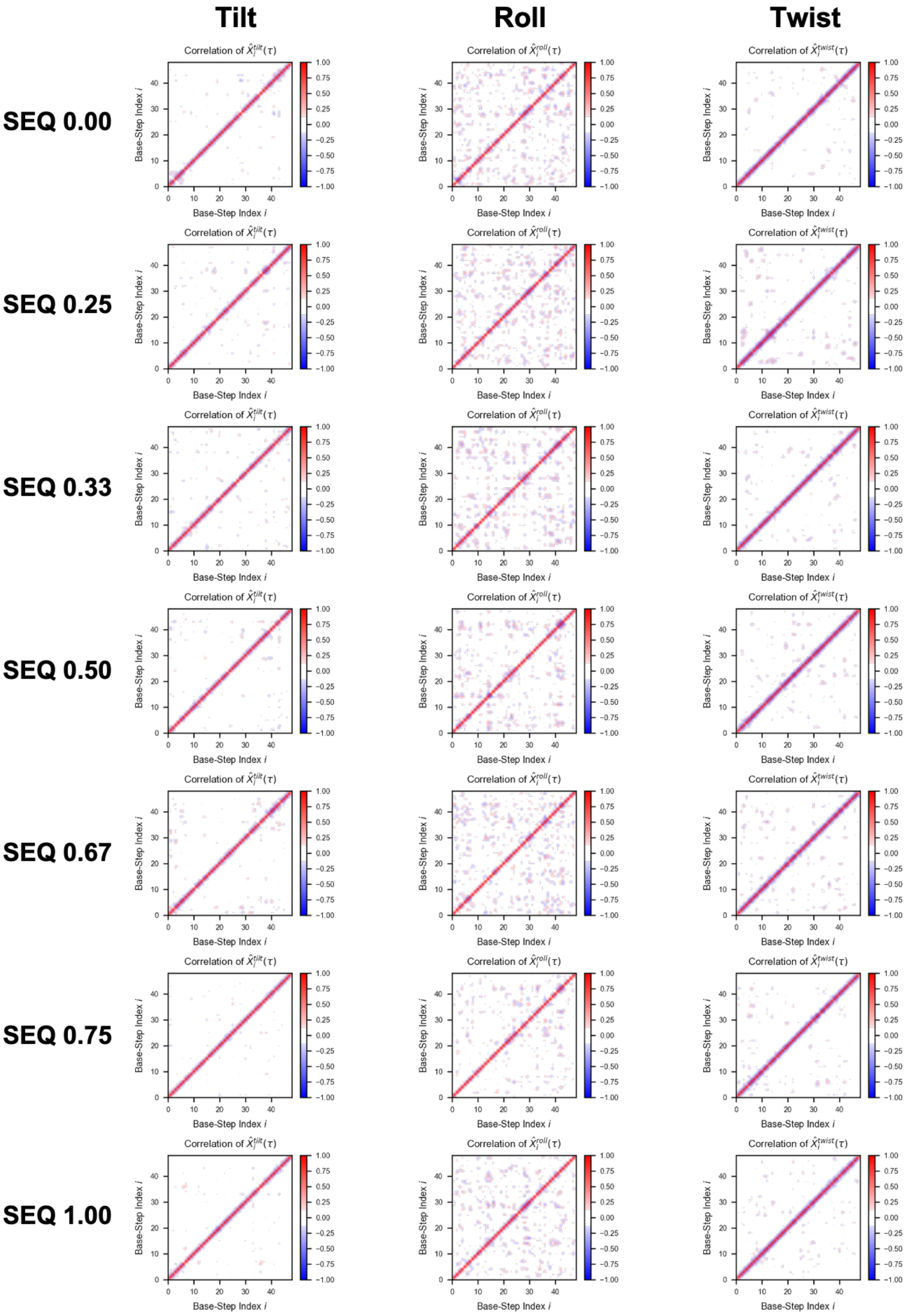

Tilt
Roll
Twist
SEQ 0.00
SEQ 0.25
SEQ 0.33
SEQ 0.50
SEQ 0.67
SEQ 0.75
SEQ 1.00
Correlation of $\hat{X}_i^{tilt}(\tau)$
Correlation of $\hat{X}_i^{roll}(\tau)$
Correlation of $\hat{X}_i^{twist}(\tau)$
Base-Step Index i
Base-Step Index j

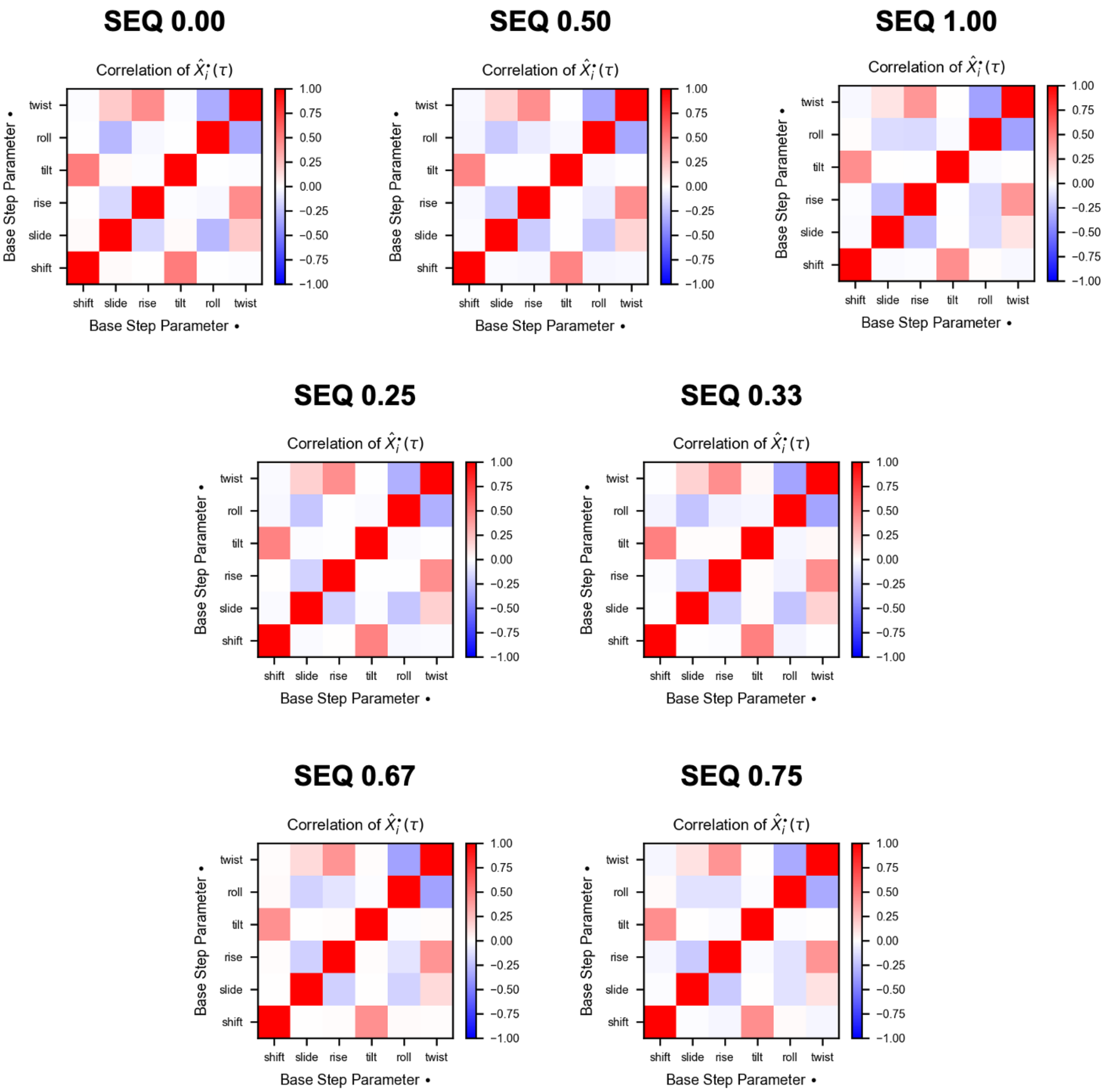


Figure S5: Correlation Coefficient Matrix of $\hat{X}_i^{\bullet}(\tau)$ between Base Step Parameters. Each matrix component is defined in Sec. II D.

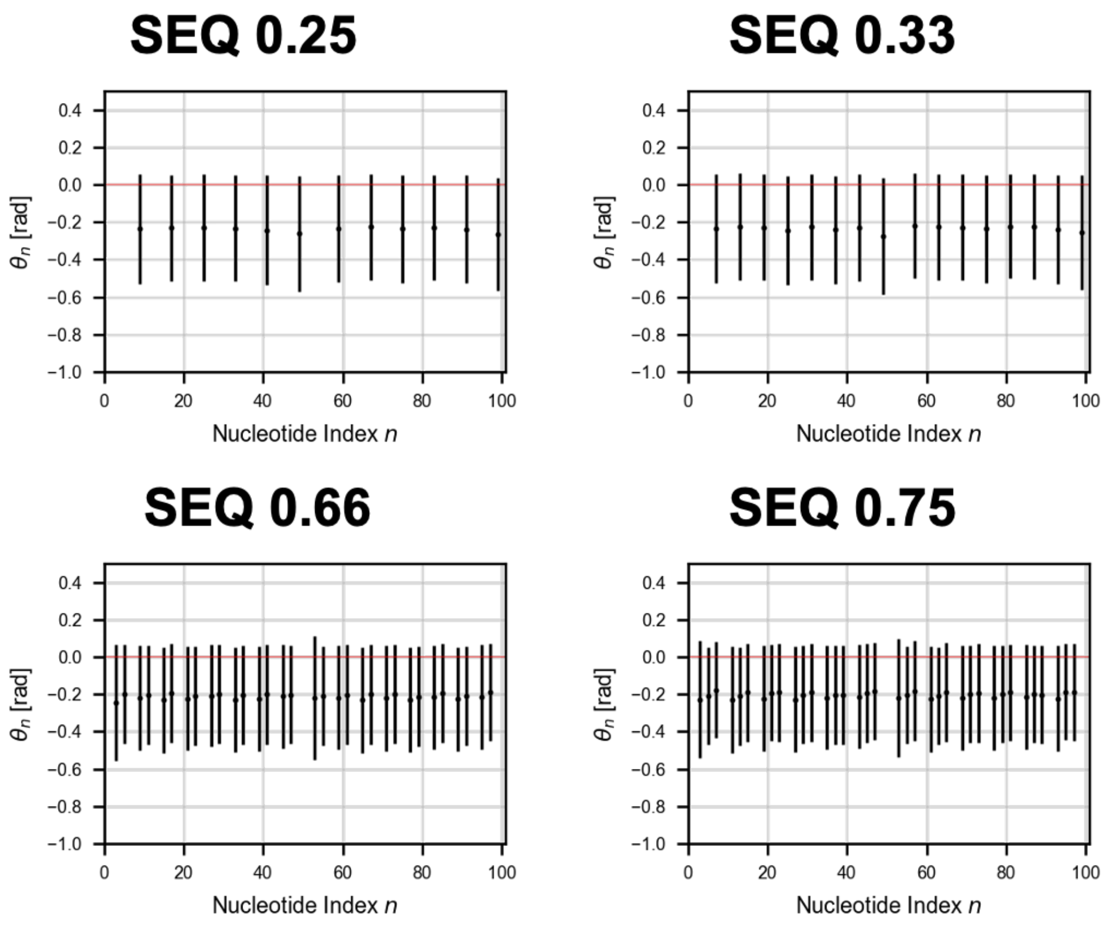


Figure S6: Orientation of Methyl Groups. The average ± the standard deviation (S.D.) of $\theta_n$ for each methyl group are shown.

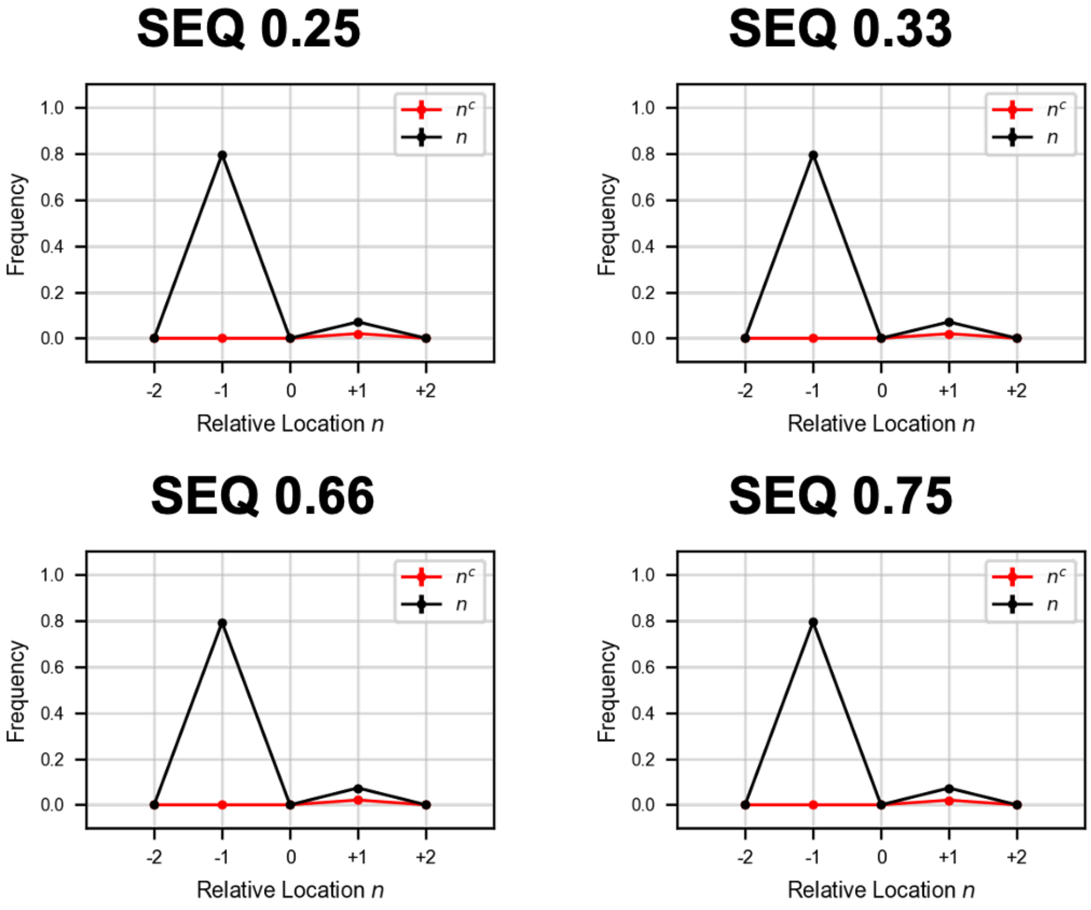


Figure S7: Interaction Between Methyl Groups and DNA. The average ± the standard deviation (S.D.) of contact frequency for all methyl groups is shown.
